## Supplementary figures and images for "A weighted constraint satisfaction approach to human goal-directed decision making"

### Supplemental Fig S1

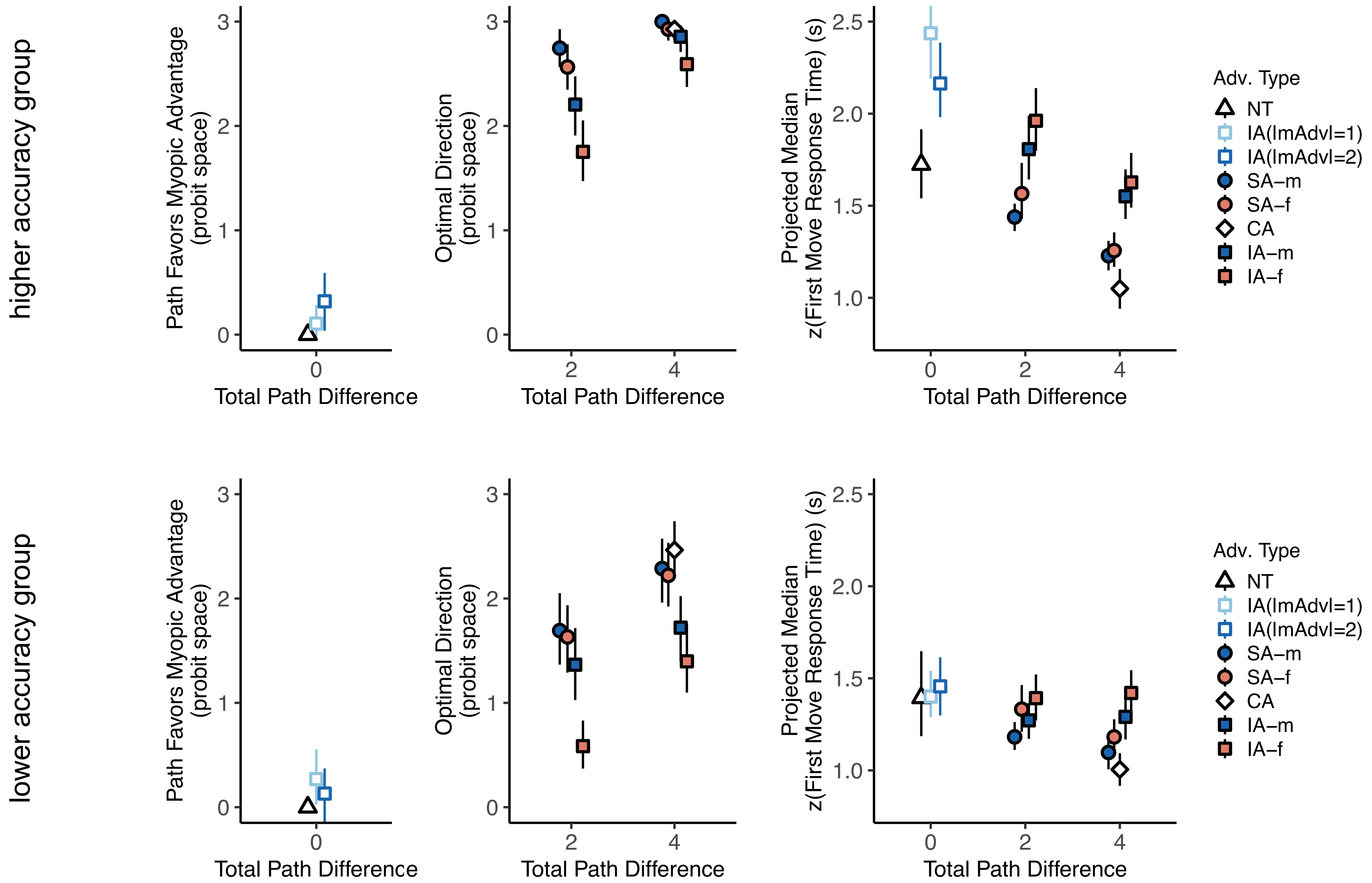

### Supplemental Fig S2

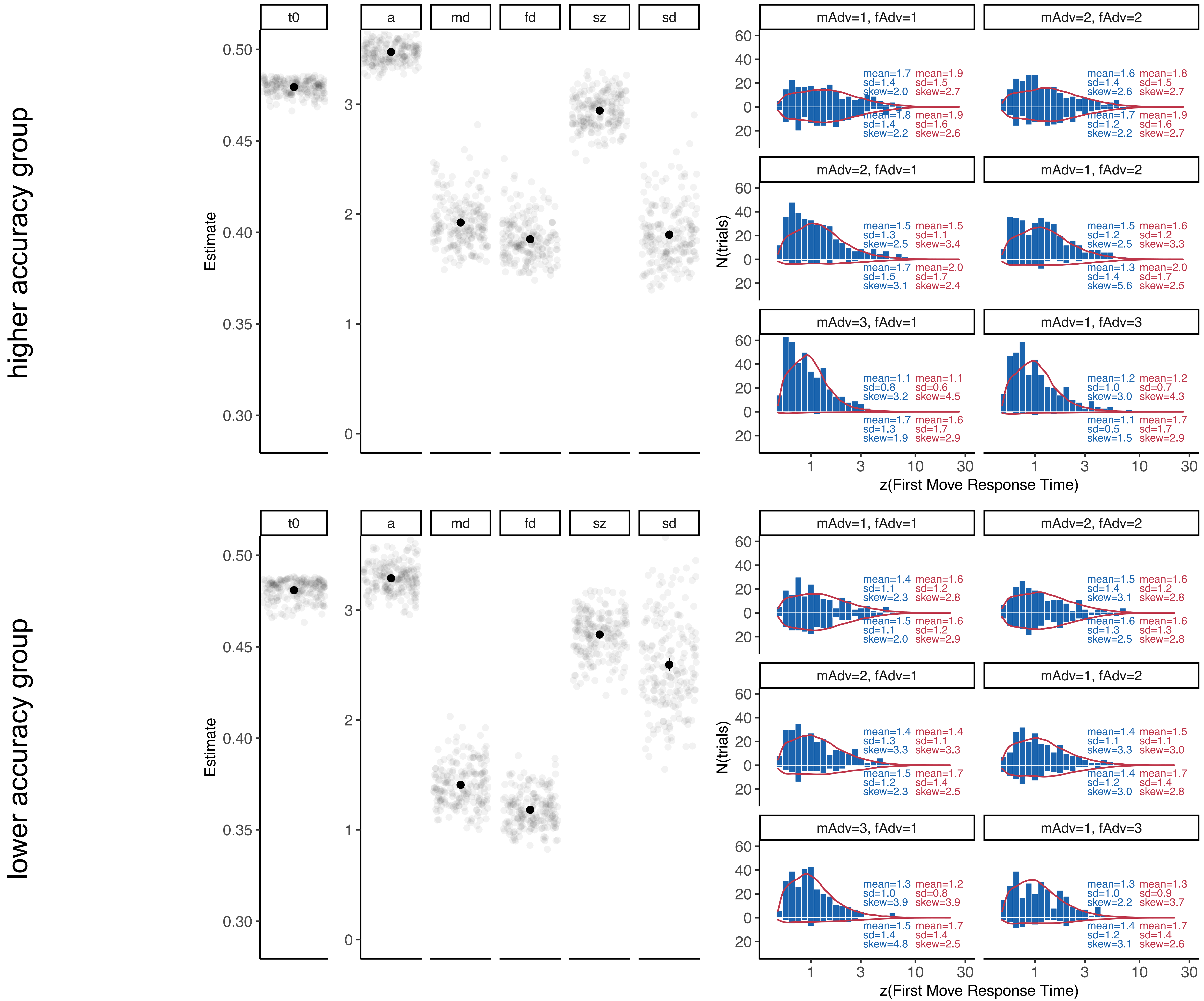

### Supplemental Fig S3

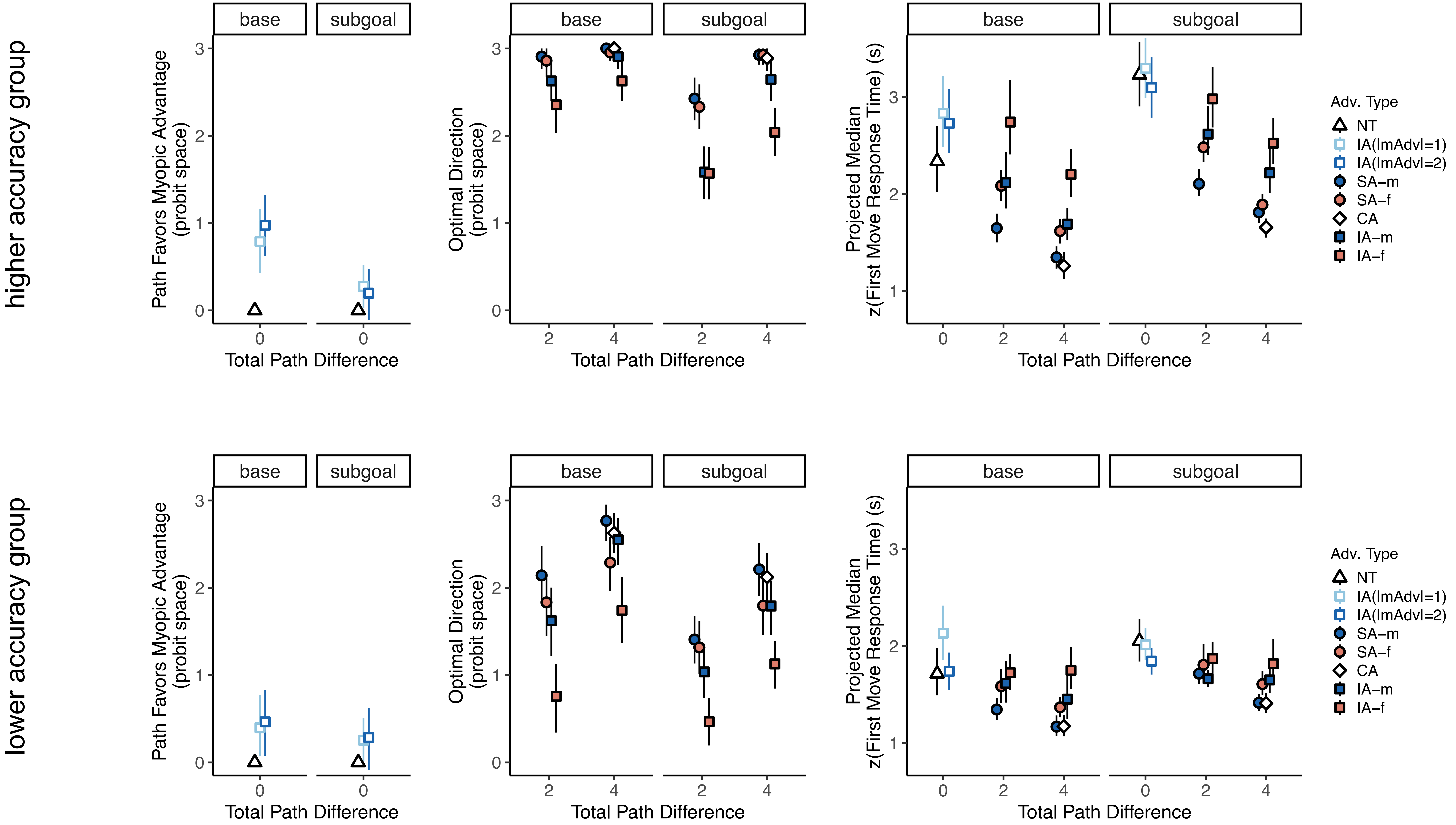

### Supplemental Fig S4

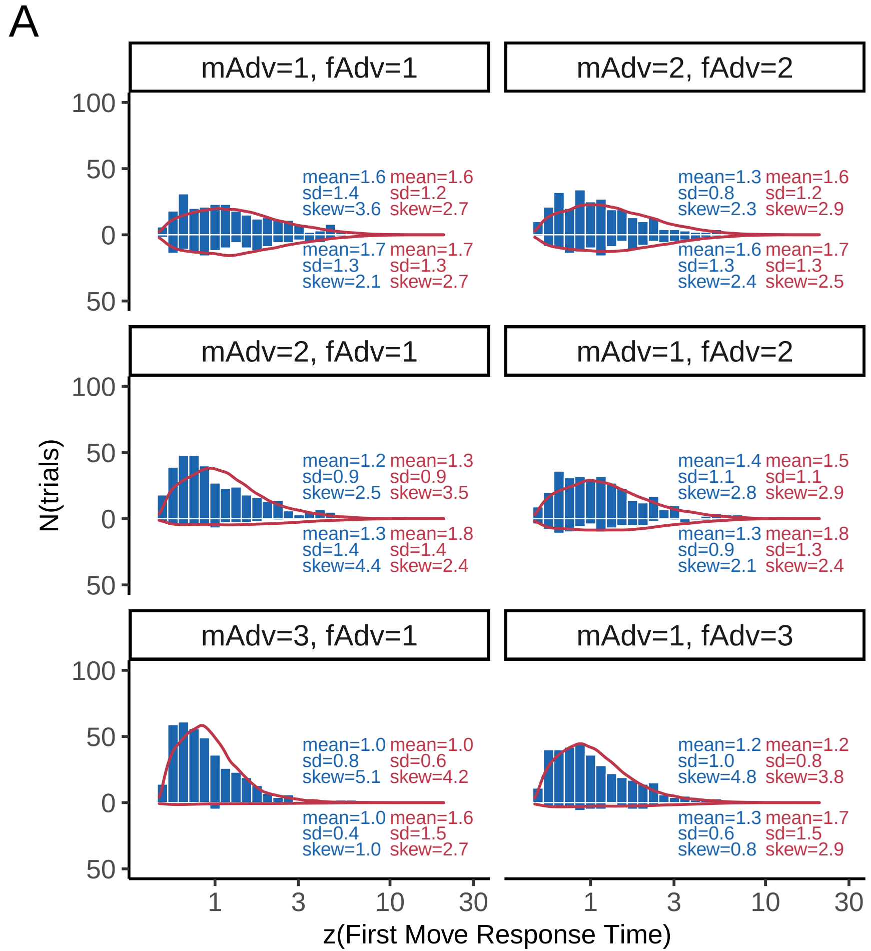

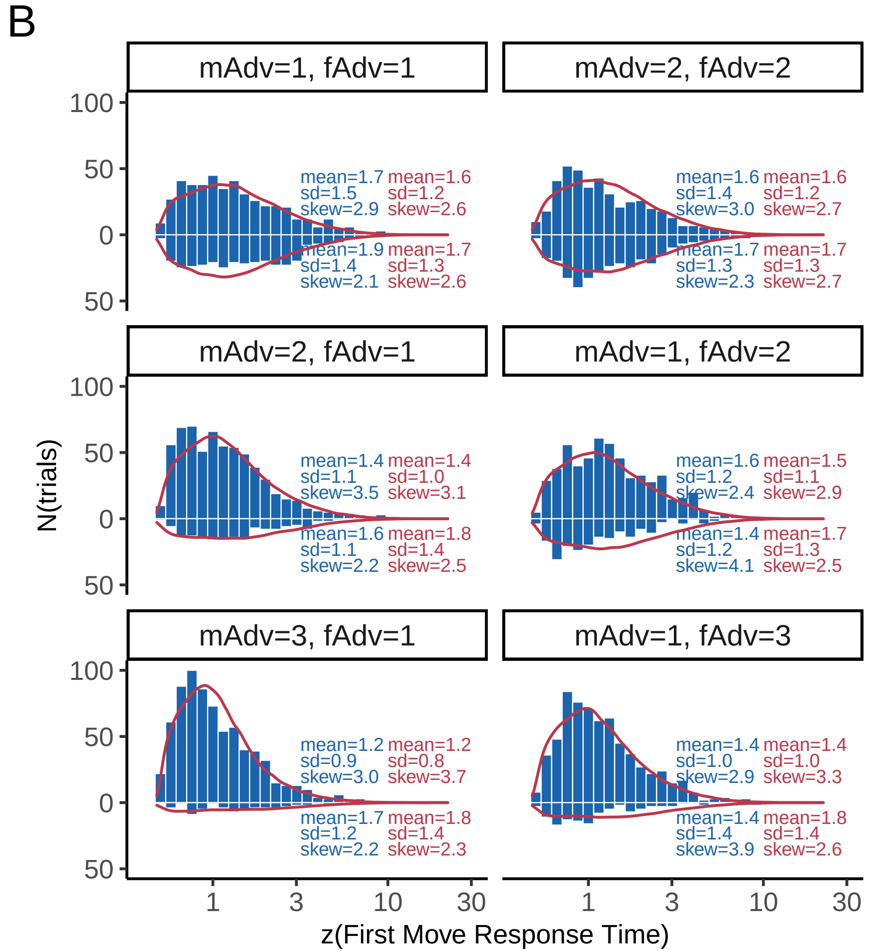

### Supplemental Fig S5

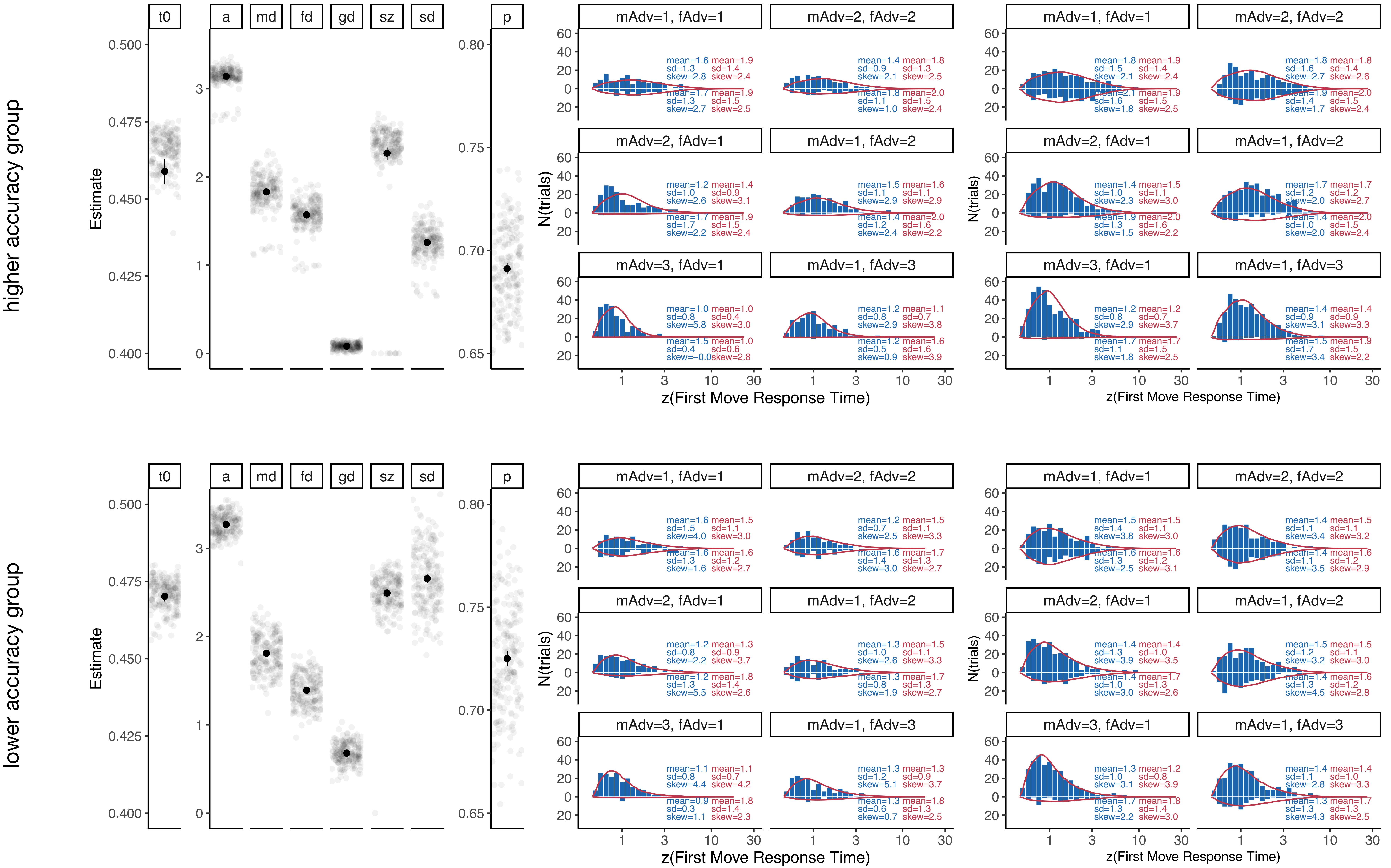

### Supplemental Fig S6

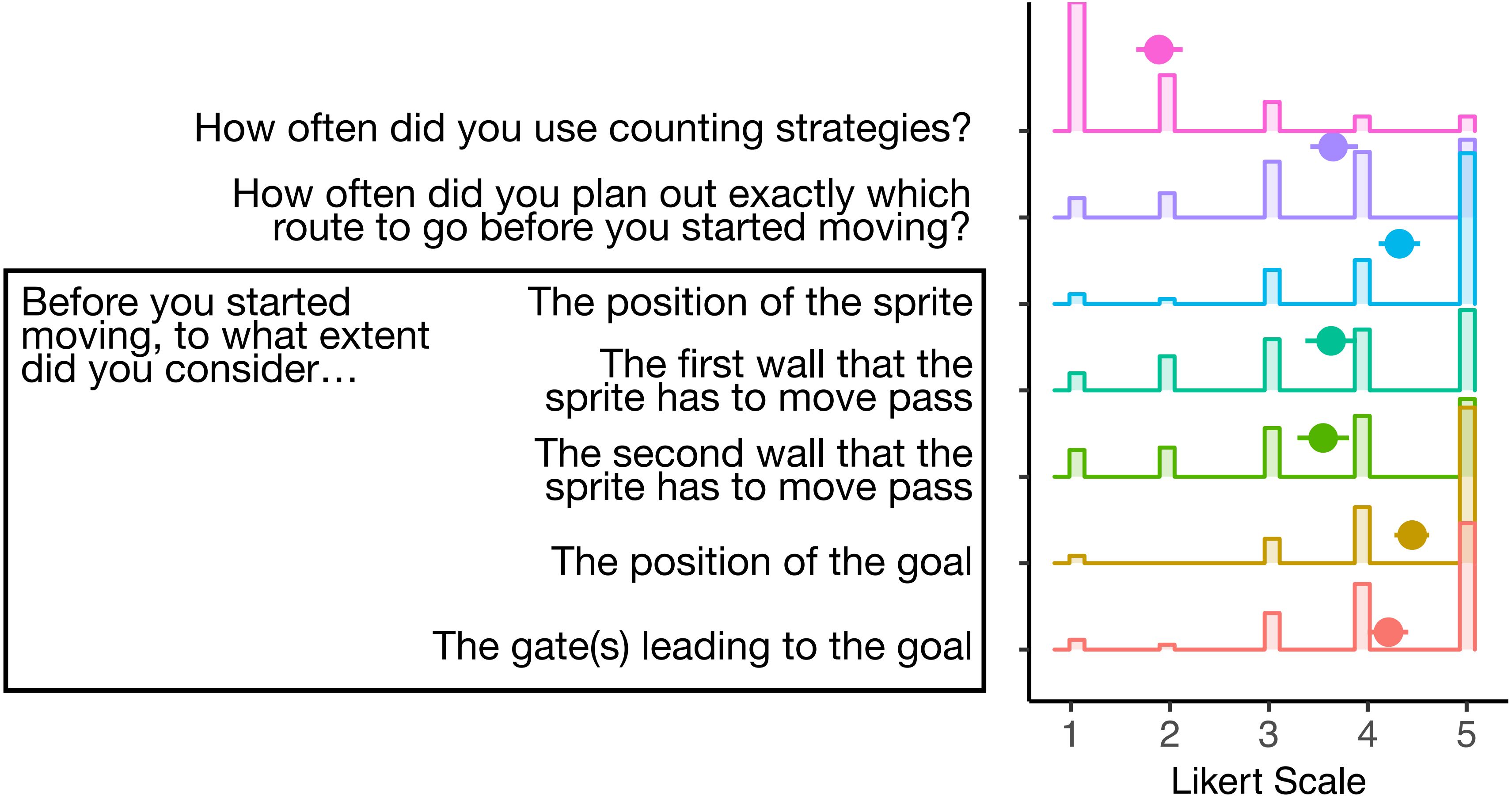
